# Self-eating while being eaten: Elucidating the relationship between aphid feeding and the plant autophagy machinery in Arabidopsis leaves

**DOI:** 10.1101/2023.03.28.534380

**Authors:** Let Kho Hao, Anuma Dangol, Reut Shavit, William Jacob Pitt, Vamsi Nalam, Yariv Brotman, Simon Michaeli, Hadas Peled-Zehavi, Vered Tzin

**Affiliations:** French Associates Institute for Agriculture and Biotechnology of Drylands, Jacob Blaustein Institutes for Desert Research, Ben-Gurion University of the Negev, Midreshet Ben Gurion, Israel; Department of Agricultural Biology, Colorado State University, CO, USA; Departments of Life Sciences, Ben-Gurion University of the Negev, Beer Sheva, Israel; Institute of Postharvest and Food Sciences, Agricultural Research Organization (ARO), Rishon LeZion, Israel; Department of Plant and Environmental Sciences, Weizmann Institute of Science, Rehovot, Israel

**Keywords:** aphid, autophagy, *Arabidopsis thaliana*, *Myzus persicae*, defense mechanism.

## Abstract

Autophagy, an intracellular process that facilitates the degradation of cytoplasmic materials, plays a dominant role in plant fitness and immunity. While autophagy was shown to be involved in plant response to fungi, bacteria, and viruses, its role in response to insect herbivory is as yet unknown. In this study, we demonstrate a role of autophagy in plant defense against herbivory using *Arabidopsis thaliana* and the green peach aphid, *Myzus persicae*. Following six hours of aphid infestation of wildtype plants, we observed high expression of the autophagy-related genes *ATG8a* and *ATG8f*, as well as *NBR1* (*Next to BRCA1 gene 1*), a selective autophagy receptor. Moreover, the number of autophagosomes detected by the overexpression of GFP-fused ATG8f in Arabidopsis increased upon aphid infestation. Following this, *atg5.1* and *atg7.2* mutants were used to study the effect of autophagy on aphid reproduction and feeding behavior. While aphid reproduction on both mutants was lower than on wildtype, feeding behavior was only affected by *atg7.2* mutants. Moreover, upon aphid feeding, the *Phytoalexin-deficient 4* (*PAD4*) defense gene was upregulated in wildtype plants but not affected in the mutants. By contrast, the hydrogen peroxide content was much higher in the mutants relative to wildtype, which might have disturbed aphid reproduction and interfered with their feeding. Additionally, an analysis of the phloem sap metabolite profile revealed that *atg7.2* mutant plants have lower levels of amino acids and sugars. These findings, together with the high hydrogen peroxide levels, suggest that aphids might exploit the plant autophagy mechanism for their survival.

## 1. Introduction

Autophagy is a well-conserved eukaryotic catabolic mechanism that is used to remove and recycle cytoplasmic components [1,2]. In plants, three distinct types of autophagy have been identified: microautophagy, macroautophagy, and megaautophagy [3,4]. Macroautophagy (hereafter referred to as autophagy) is well-characterized in plants and other organisms [5]. Its pathway is characterized by the formation of double-membrane vesicles, named autophagosomes, that sequester cytosolic components such as specific proteins, protein aggregates, damaged organelles, or organelle components, and carry them to the vacuole for degradation [6]. The genes functioning in the autophagy machinery, autophagy-related (*ATG)* genes, were first discovered through forward-genetic screens for autophagy-defective mutants in yeast (*Saccharomyces cerevisiae*) and are highly conserved [7–9]. Over the past few decades, more than 40 conserved *ATGs* have been identified in yeast, animals, and plants [3]. Nearly half of the identified *ATG* genes are part of the core autophagy machinery that is conserved across kingdoms, including in Arabidopsis [10].

Expression studies, as well as the combined use of *ATG* knock-out mutants such as ATG and ATG7, and autophagy markers such as ATG8 brought to light the important roles of autophagy in plant homeostasis and adaptation to environmental stresses [11–13]. Autophagy has been shown to function in plants in response to various abiotic stresses such as starvation [14], high salinity [15], drought [16], heat [17], chilling stress [18], and hypoxia [19], most of which lead to osmotic or oxidative stresses [20]. Autophagy induction in response to these stresses can assist in nutrient recycling and mobilization, as well as removal of oxidatively damaged proteins and organelles. The role of autophagy in plant biotic stress responses has been studied mainly in relation to infection with pathogens such as fungi, bacteria, and viruses [21–23]. Autophagy activation can lead to different outcomes depending on the lifestyle of the pathogen or the pathosystem, and autophagy was shown to have both pro-survival and pro-death activities. For instance, autophagy was shown to play an antiviral role in plant-virus interactions, but increasing evidence suggests that viruses can also exploit the autophagy pathway to promote pathogenesis [21].

Insect herbivory represents a major challenge to plants’ growth. Hence, plants have developed an array of mechanisms to protect themselves from herbivorous insect attacks, such as activating different metabolic pathways, which considerably alter their chemical and physical properties [24]. For instance, central and specialized metabolism are modified, the photosynthetic efficiency is either elevated or suppressed, and nutrients such as carbon and nitrogen are remobilized [25,26]. The metabolic adjustment can affect phloem quality and metabolite composition [27], directly affecting phloem sap-feeding insects [27,28]. Moreover, the production of defensive compounds requires a high amount of energy, which causes a significant demand for resources [26,29]. Plants cope with this challenge by degrading or remobilizing resources such as carbohydrates and proteins, to keep up with the required energy demand [25,29,30]. Though these processes bring to mind the autophagy machinery, the only evidence for autophagy involvement in plant defense mechanisms against insect herbivores is the induction of several *ATG* genes by *Myzus persicae* (green peach aphid; GPA) infestation [31–33]. Thus, the role of the plants’ autophagy machinery in responses to insect herbivores has yet to be fully revealed.

Here, we investigated the relationship between insect infestation and the autophagy machinery in plants by focusing on two well-studied model organisms, *Arabidopsis thaliana* and GPA. This compatible pathosystem has been successfully utilized to characterize plant responses against phloem-feeding insects and to identify plant genes and mechanisms contributing to defense against phloem sap-feeding insects [34–36]. Using a variety of experimental approaches, including gene expression analysis, autophagosome formation, insect bioassays, metabolic profiling, and detection of hydrogen peroxide, this study aims to elucidate the possible interaction between the autophagy machinery and insect herbivore infestation in plants.

## 2. Results

### 2.1. Aphids infestation induces expression of ATG genes and increases the number of autophagosomes

To determine whether *ATGs* are induced in response to aphid feeding, wildtype plants were infested with GPA for 6 h. The expression levels of four autophagy genes, *ATG5, ATG7, ATG8a, and ATG8f,* were measured, as well as the selective autophagy receptor gene *NBR1* (*Next to BRCA1 gene 1*). Gene expression levels were normalized to the reference gene *PP2A* and presented as fold change relative to the untreated control. As shown in Figure 1, the *ATG8a, ATG8f,* and *NBR1* genes were significantly upregulated upon aphid feeding, while *ATG5* and *ATG7* were not affected.

**Figure 1.**
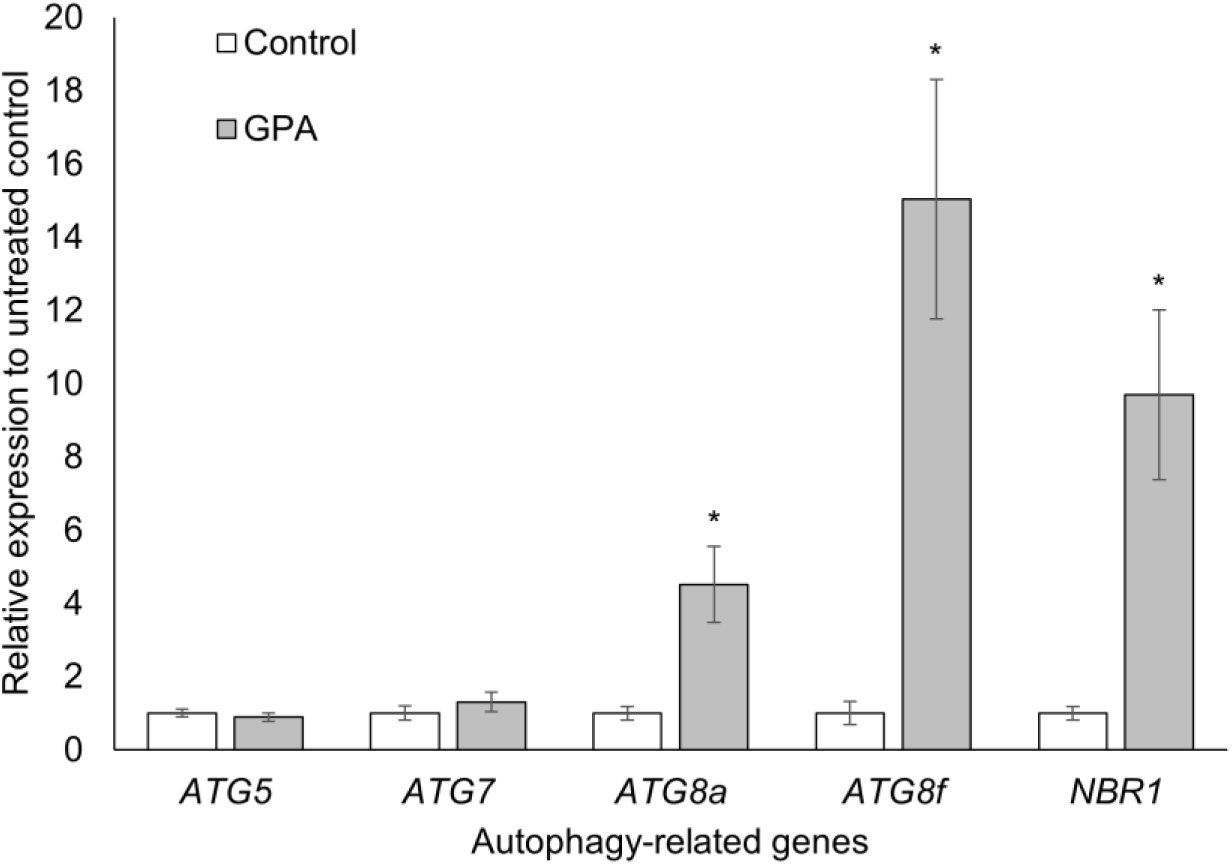
The effect of aphid feeding on the expression levels of autophagy-related genes. Leaves of Col-0 wildtype plants were infested with GPA or left untreated (Control). The expression levels of five autophagy-related genes were quantified using qRT-PCR and normalized to a reference gene, *PP2A*. The values are presented in fold change relative to the control of each gene. Asterisks indicate statistical significance * *P* < 0.05, Student’s *t*-test. Error bars indicate standard errors of the mean (n = 3-4).

ATG8, which in plants exists as a gene family, is a core component of the autophagy machinery. It is synthesized as a proprotein and goes through several processing events that result in its covalent attachment to phosphatidylethanolamine (PE) at the autophagosomal membrane. As it is found on the autophagosome from its formation to its lytic destruction in the vacuole, a fluorescently tagged ATG8 is commonly used as an autophagosome marker [37]. To look at autophagy induction in response to aphid infestation, leaves of an Arabidopsis line that expresses GFP-ATG8f were infested with GPAs and GFP-labeled autophagosomes were detected by confocal fluorescence microscopy. Concanamycin-A, an inhibitor of vacuolar H+-ATPase, was used to increase vacuolar pH and inhibit vacuolar enzymes activity. Under these conditions, autophagic bodies accumulate in the vacuole and there is an increase in the amount of autophagosomes in the cytoplasm, facilitating the visualization of autophagy processes [38,39]. As shown in Figure 2, the number of fluorescently labeled puncta in GPA-treated leaves was approximately three times higher than in the control leaves. No effect of concanamycin-A was observed relative to the control (Figure 2B). Altogether, the gene expression and autophagosome formation results suggest that aphid infestation induced the autophagy machinery in Arabidopsis leaves.

**Figure 2.**
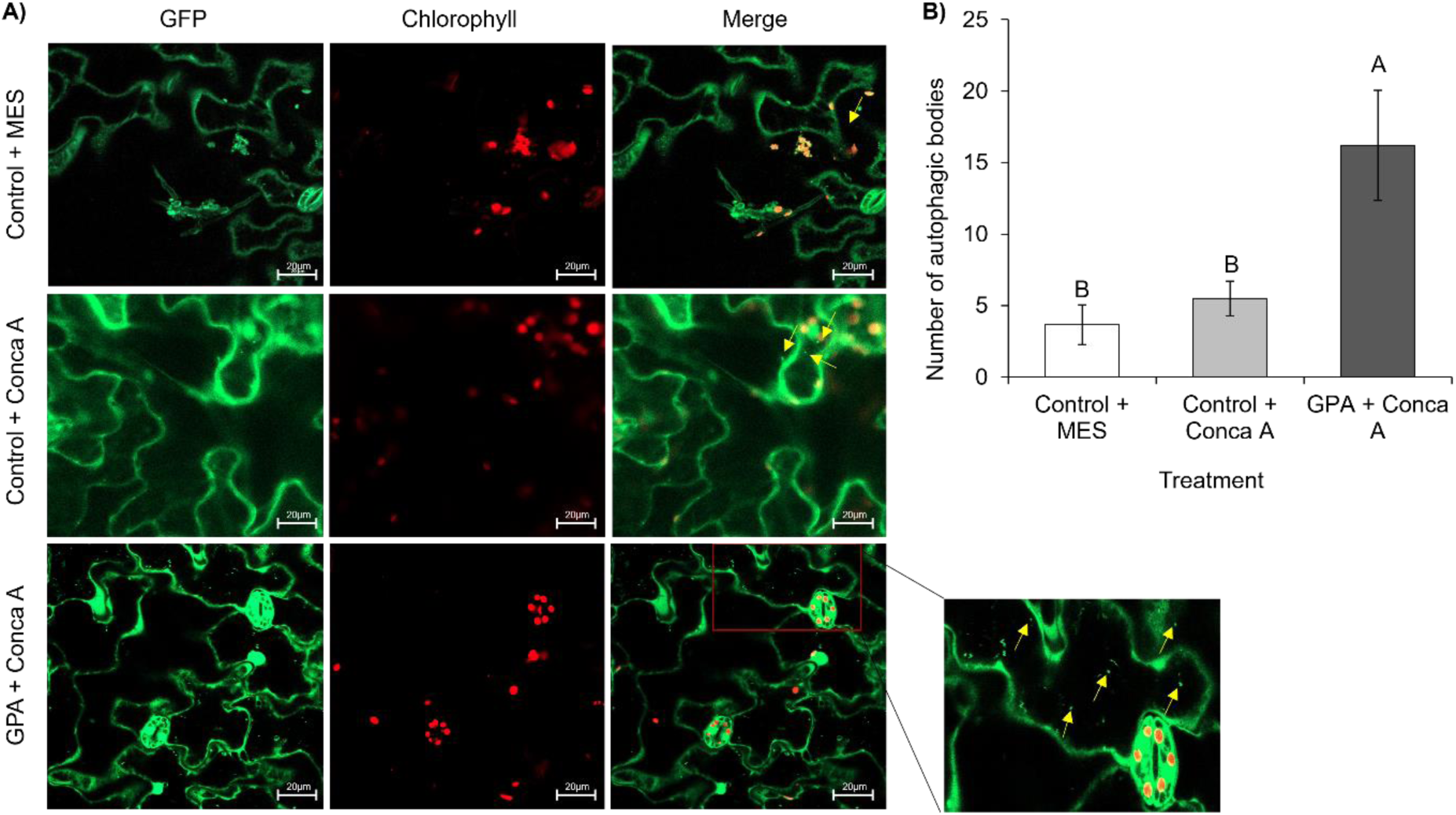
Autophagy activation in response to aphid infestation. Leaves of GFP-ATG8f transgenic plants were infested with 20 adult GPAs for 72 h and visualized under a confocal microscope to determine whether aphid feeding induced autophagy. (A) Representative confocal images of GFP-ATG8f transgenic leaf grown under normal growth conditions or aphid infestation with or without the addition of concanamycin-A (Conca A). Yellow arrows indicate GFP-ATG8f labeled puncta. (B) Quantification of autophagic bodies in GFP-ATG8f transgenic leaves. The average number of autophagic bodies was calculated for each condition, and statistical significance was determined using one-way analysis of variance. Different letter codes indicate significant differences in concentrations at *P* < 0.05, as indicated by one-way ANOVA with post hoc Tukey’s analysis. Error bars indicate standard errors of the mean (n = 6).

### 2.2. *Autophagy-deficient* mutants affect aphid performance and feeding behavior

Two *autophagy-deficient* mutants, *atg5.1* [40], and *atg7.2* [41] were used to determine whether the autophagy machinery affects GPA feeding and behavior. These T-DNA insertion knockouts are extensively used for studying the autophagy machinery in plants [42,43]. Reduction in the expression levels of *ATG5* and *ATG7* in the mutants was verified by qRT-PCR (Figure S1). Then, a no-choice bioassay was conducted to measure changes in GPA body weight and reproduction. As shown in Figure 3A, the weight of the GPAs that fed on *atg* mutant plants was significantly lower than on the wildtype. To test the effect on aphid fecundity, the number of total aphids (nymphs and adults) was evaluated after seven days of infestation. The results showed that GPAs reproduce less well on the two *atg*-deficient mutants compared to wildtype (Figure 3B). The reduction in body weight and reproduction of the GPAs might be due to either a poor diet and/or differential induction of plant defense mechanisms in the autophagy-deficient plants.

**Figure 3.**
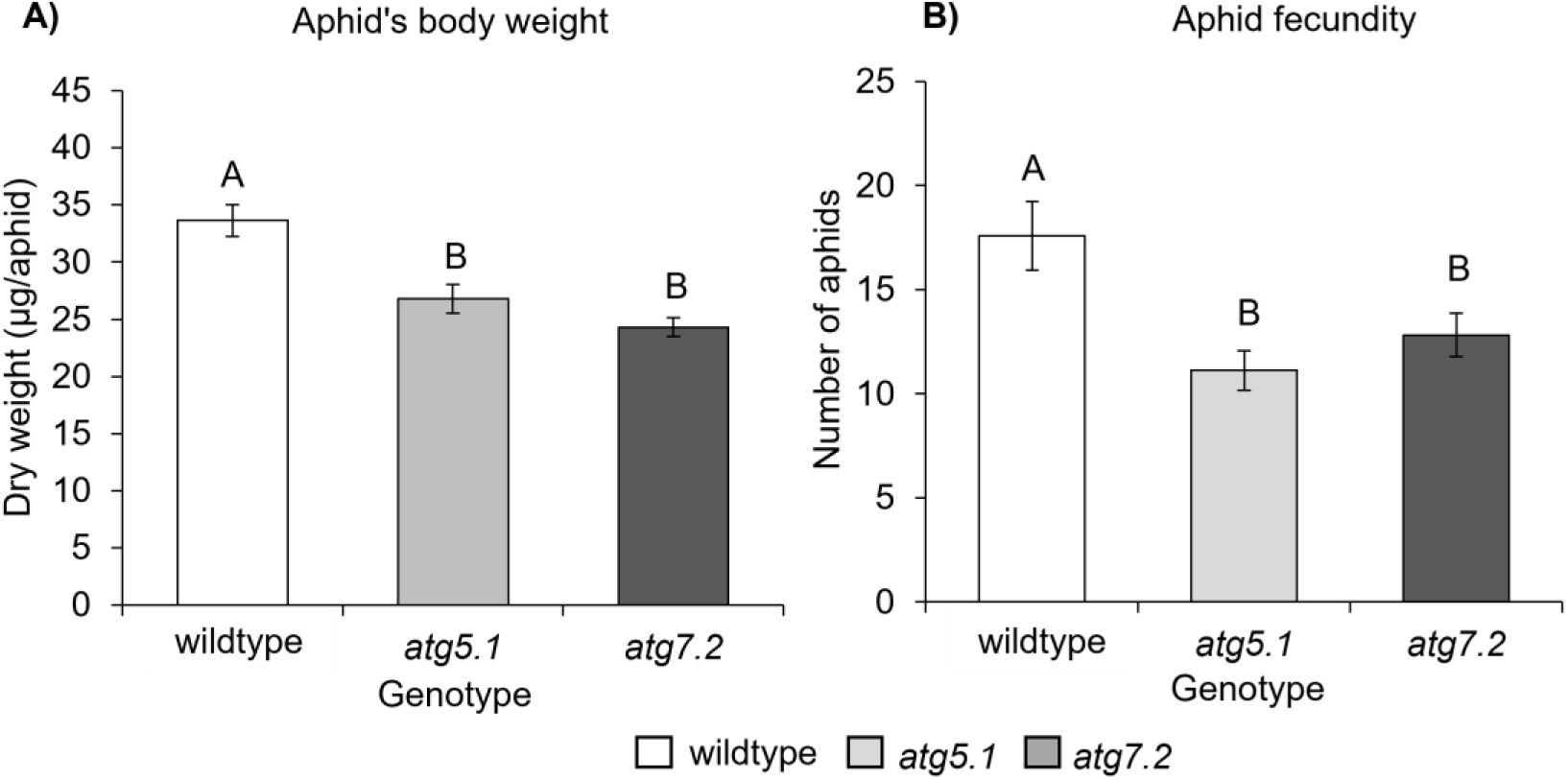
The effect of autophagy-deficient mutants on aphid growth and reproduction. (A) Aphid body weight was measured following 6 h of feeding on *atg* mutants or wildtype plants (n = 4). (B) Aphid fecundity was compared after 7 d of infestation by counting the total number of nymphs and adults (n = 12). Different letter codes indicate significant differences at *P* < 0.05, as indicated by one-way ANOVA with post hoc Tukey’s analysis. Error bars indicate standard errors of the mean.

To further characterize the effect of autophagy on GPA physiology, their feeding behavior was evaluated using an Electrical Penetration Graph (EPG) assay. This assay measures the electromotive force signal and fluctuations in electrical resistance resulting from aphid stylet penetrations, and is commonly used to monitor the feeding behavior of phloem feeders across leaf tissues (i.e., phloem, xylem, epidermis, or mesophyll) and penetration through the leaf surface [44]. The effect of autophagy on GPA feeding behavior was compared by analyzing the parameters from the four main EPG phases. The results showed that GPA feeding behavior was significantly different between the *atg7.2* mutant and wildtype plants, while no effect was detected in the *atg5.1* mutant (Table 1). The occurrence of events of GPA feeding in the phloem (n_E2) and the time spent in phloem ingestion (%probtimeinE2) were significantly lower, and the duration of aphid probing of the epidermis and mesophyll tissues (%probtimeinC) was longer when fed on *atg7.2* mutant compared to wildtype (Supplementary Table S2). Taken together, our results suggest that autophagy deficiency in Arabidopsis plants affects aphid body weight, fecundity, and feeding behavior.

**Table 1.**
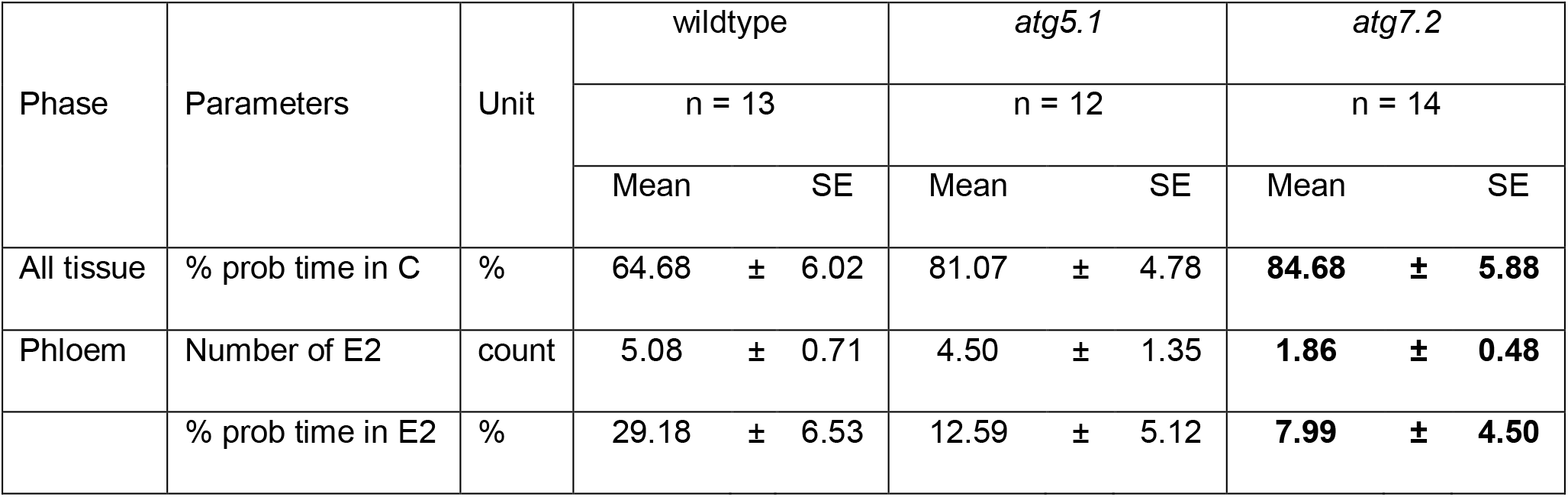
Feeding behavior of GPAs on *atg* mutants. Waveforms were analyzed using Stylet^+^a software, and an Excel workbook for automatic parameter calculation [45]. In bold are significant parameters relative to wildtype (Wilcoxon test, Adj. *P* < 0.05).

| Phase | Parameters | Unit | wildtype |  | *atg5.1* |  | *atg7.2* |  |
| --- | --- | --- | --- | --- | --- | --- | --- | --- |
|  |  |  | n = 13 |  | n = 12 |  | n = 14 |  |
|  |  |  | Mean | SE | Mean | SE | Mean | SE |
| All tissue | % prob time in C | % | 64.68 | ± 6.02 | 81.07 | ± 4.78 | **84.68** | **± 5.88** |
| Phloem | Number of E2 | count | 5.08 | ± 0.71 | 4.50 | ± 1.35 | **1.86** | **± 0.48** |
|  | % prob time in E2 | % | 29.18 | ± 6.53 | 12.59 | ± 5.12 | **7.99** | **± 4.50** |

### 2.3. Metabolic profile of autophagy-deficient mutants upon aphid infestation

The aphid no-choice bioassays and EPG analysis suggested that aphids possess different feeding behaviors on the *atg* mutants compared to wildtype. This might be due to a decreased attractiveness to insects in terms of nutrient composition in the phloem sap and/or due to difference in the defense responses. To explore the effect of metabolite composition, we performed a GC-MS analysis measuring the central metabolites in the phloem sap of *atg* mutants. Overall, 33 compounds were detected in the phloem sap of GPA-treated and untreated leaves of wildtype and *atg* mutants (Supplementary Table S3). First, the metabolites were clustered using hierarchical clustering with Euclidean distance measure and ward agglomeration method and visualized in a heatmap to get an overview of metabolite patterns by genotype and GPA treatments (Figure 4A). Without GPA treatment, *atg7.2* and wildtype were clustered together, separated from *atg5.1*. Upon GPA infestation, a large metabolic difference was observed in GPA-infested *atg5.1* mutant and wildtype plants, compared to uninfested plants. By contrast, *atg7.2* was closer to uninfested *atg7.2* and wildtype. Similar modification of metabolic profiles in the phloem of wildtype and *atg5.1* was observed under GPA feeding, which is in accordance with the EPG results (Table 1).

**Figure 4.**
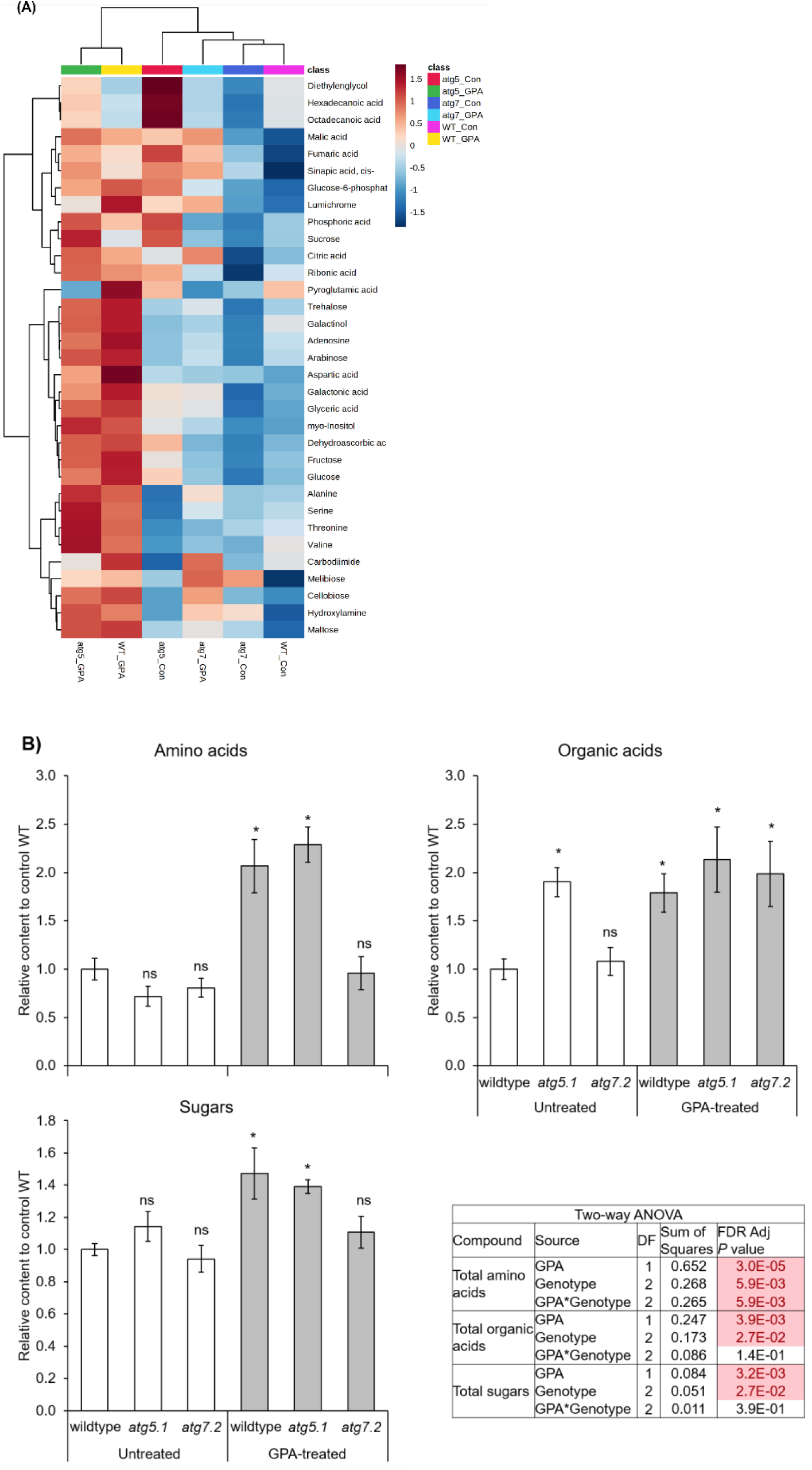
A targeted metabolic overview of *atg* mutants infested with aphids for 6 h. (A) A heatmap analysis, presenting central metabolites profile. The Euclidean distance with Ward’s minimum variance method was calculated using the default parameters of the MetaboAnalyst software. Colors correspond with concentration values (autoscale parameters), where red indicates high levels and blue indicates low levels. (B) Relative levels of total organic acids, amino acids, and sugars in the phloem of untreated or GPA treated wildtype and *atg* mutant plants. Metabolite content is shown relative to untreated wildtype plants. Asterisks indicate statistical significance * *P* < 0.05, Dunnett’s test. ns, not significant. Error bars indicate standard errors of the mean, n = 5.

Next, we performed a two-way ANOVA analysis to identify significantly altered metabolites. The levels of 23 metabolites were significantly affected by either genotype, GPA treatment or their interaction (Supplementary Table S4). High levels of GPA-induced organic acids were observed in the wildtype and *atg7.2* mutant, while *atg5.1* showed higher organic acids levels at basal but no increased upon GPA infestation. Exposure to aphids caused an accumulation of amino acids and sugars in the *atg5.1* mutant and wildtype, but not in the *atg7.2* mutant. Overall, the total amino acid content was significantly affected by GPA treatment, genotype, and genotype/GPA treatment interaction, while total organic acids and total sugars were significantly affected by genotype and GPA treatment but not their interaction (Figure 4B). Among these metabolites, three amino acids (serine, threonine, and valine) and one organic acid (fumaric acid) were affected by genotype, GPA treatment, as well as their interaction (Supplementary Table S4). Upon GPA treatment, these amino acids were highly induced in *atg5.1* plants, while only fumaric acid was induced in *atg7.2* plants. In wildtype plants, increased levels of serine and fumaric acid were observed under GPA infestation (Figure 5). In addition, the basal levels of serine and fumaric acid in *atg5.1* were lower and higher, respectively, compared to untreated control. Overall, the metabolic analysis suggests *atg5.1* showed similar response as wildtype to GPA feeding, while *atg7.2* has a different pattern. This correlates with the EPG results, suggesting that the difference in feeding behavior might be the result of different nutrient content.

**Figure 5.**
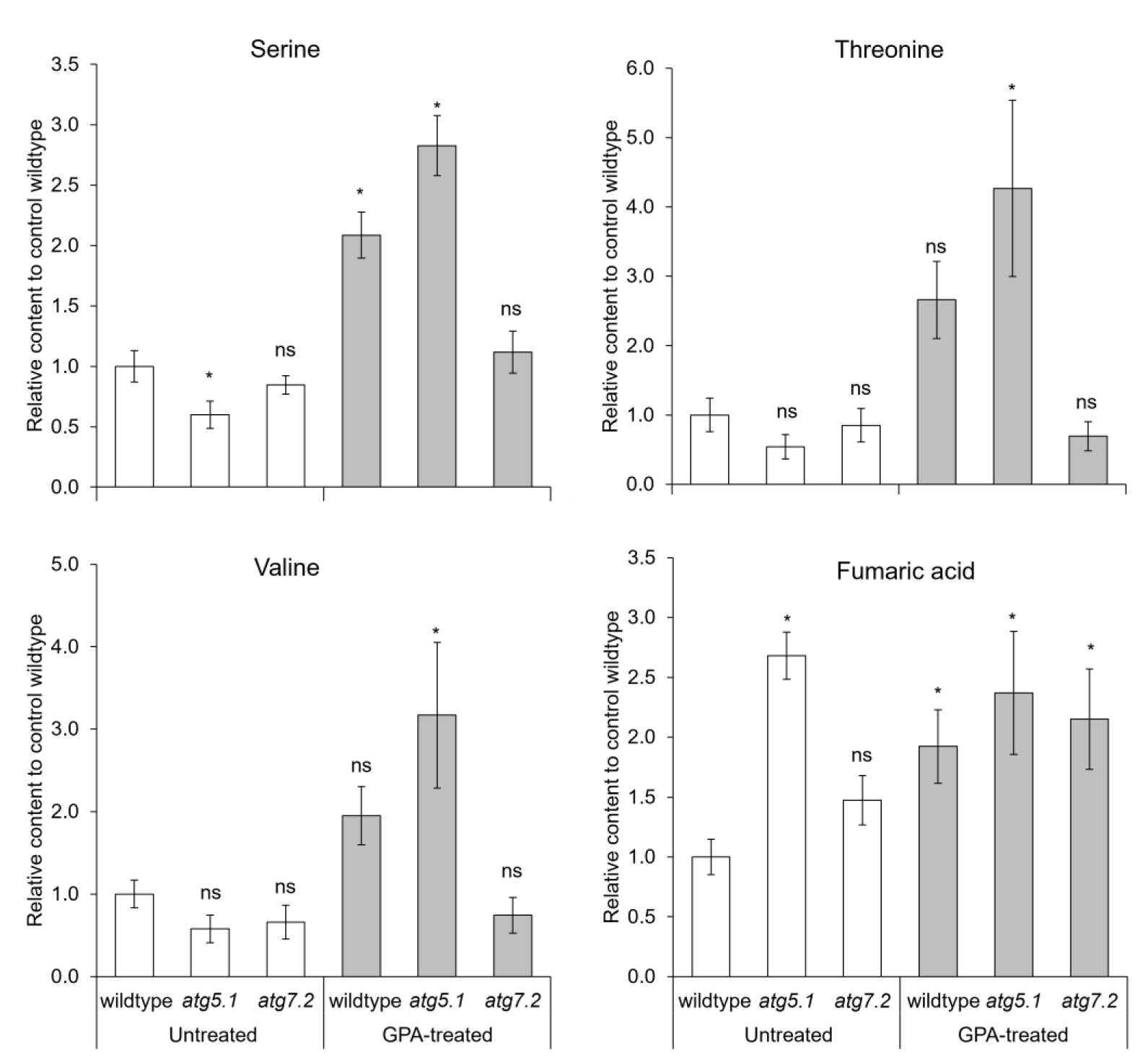
The effect of GPA feeding on the metabolic profile of the phloem sap of *atg* mutants and wildtype plants. Relative levels of serine, threonine, valine and fumaric acid in the phloem sap of Arabidopsis *atg* mutants and wildtype plants with or without aphid treatment, compared to untreated wildtype plants. Asterisk indicates significant differences in concentrations at *P* <0.05 level indicated by Dunnett’s Student’s *t*-test, and n.s. stands for not significant. Error bars indicate standard errors of the mean, n = 5.

### 2.4. The effect of aphid feeding on the defense mechanism of *atg* deficient mutants

To test the hypothesis that differential activation of defense mechanisms in the *atg* mutant plays a role in the reduced body weight and fecundity of GPAs feeding them, we measured the expression level of *Phytoalexin deficient 4* (*PAD4*). *PAD4* is a defense-related gene that is involved in stimulating the production of the defense phytohormone salicylic acid (SA), as well as other processes that limit pathogen and aphid growth [46–49]. As presented in Figure 6, the expression of *PAD4* in wildtype plants was significantly increased upon GPA feeding, while it was not affected in both *atg* mutants. This suggests that the reduction of aphid performance on the *atg* mutants (Figure 3, and Table 1) is not the result of the induction of the plants defense response *via* SA signaling. However, it is possible that it is the result of activation of other defense mechanisms.

**Figure 6.**
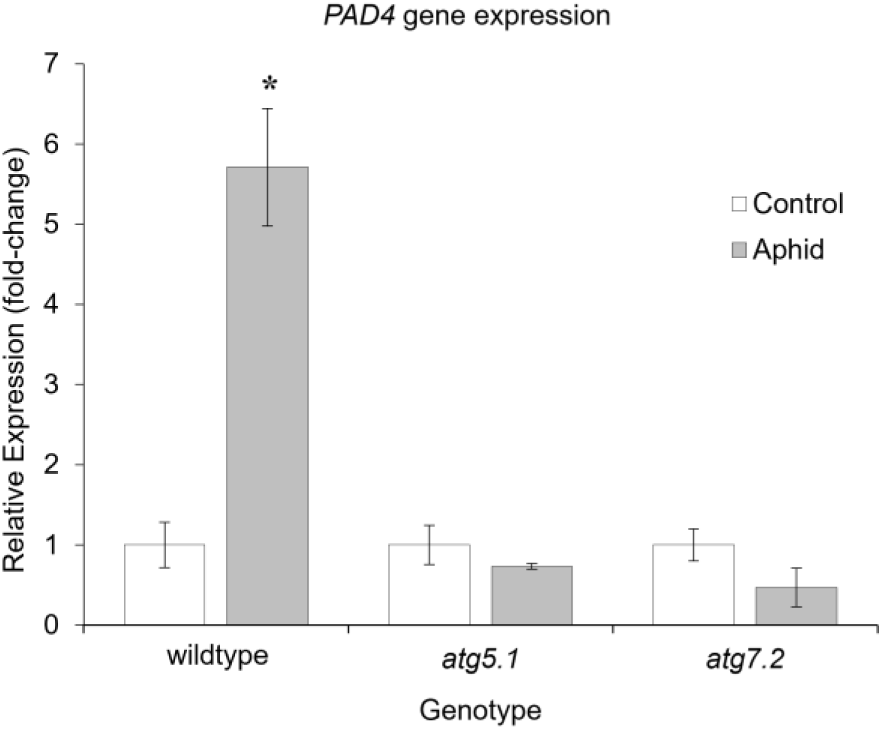
*Phytoalexin deficient 4* (*PAD4*) gene expression. *PAD4* gene expression was measured in GPA infested and control leaves of wildtype and *atg* mutants along with a reference gene, *PP2A*. The values are presented as fold change relative to the control of each genotype. Asterisks indicate statistical significance * *P* < 0.05, Student’s *t*-test. Error bars indicate standard errors of the mean (n = 4).

Thus, we conducted DAB staining to detect the presence of hydrogen peroxide, the most stable type of reactive oxygen species (ROS). ROS are involved in signaling cascades in response to many environmental stresses, and are known to be involved in plant defense against aphids [50]. Under the control condition (without aphids), the hydrogen peroxide levels were high in both *atg5.1* and *atg7.2* relative to the wildtype, and did not change much upon aphid infestation. Hydrogen peroxide levels increased in the wildtype plants following aphid infestation but did not reach the levels observed in the *atg* mutants (Figure 7). The high levels of hydrogen peroxide observed in the *atg*-deficient mutants might explain the poor GPA performance and feeding behavior (Figure 3).

**Figure 7.**
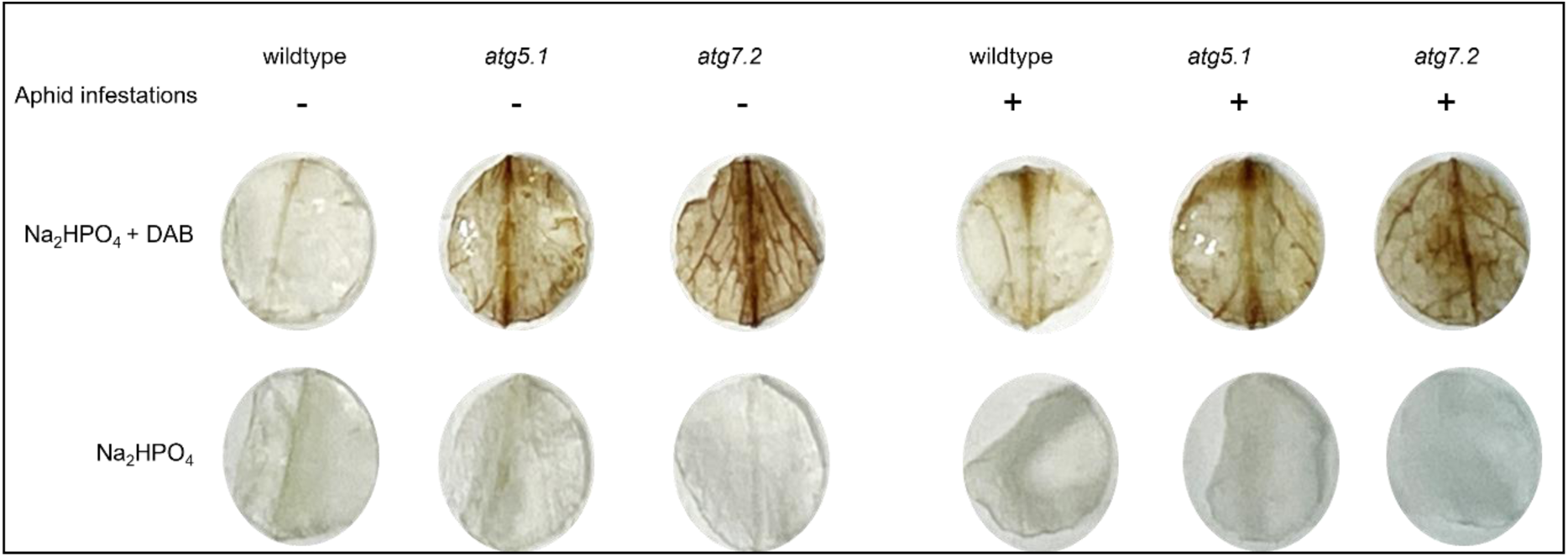
Physiological characterization of hydrogen peroxide levels in *atg* mutant leaves using DAB staining. The measurements were conducted under GPA treated and untreated conditions (7 d). Upper panel: Na_2_HPO_4_ + DAB solution; lower panel: Na_2_HPO_4_ solution, which was applied as a control treatment.

## 3. Discussion

### 3.1. Aphids affect the autophagy machinery

Our research highlights as yet unfamiliar relationship between autophagy machinery and insect herbivory. We investigated whether phloem-feeding insects induce autophagy in Arabidopsis plants and their potential interactions. Previous studies showed significant upregulation of *ATG* genes and proteins under various biotic or abiotic stresses [51–55], but only a few studies aim to reveal this relationship between plants and insects. A study from 2006 by Seay *et al*. used the microarray data available on GENEVESTIGATOR database and suggested that Arabidopsis plants infested with GPA showed that autophagy-related genes *ATG4*, *ATG8,* and *ATG18* were significantly induced upon aphid infestation [31]. In addition, Kuśnierczyk *et al.,* 2007 showed an induction of *ATG8a*, *ATG8e*, *ATG8f*, and *NBR1* in Arabidopsis Wassilewskija ecotype after 72 h feeding of GPA [33], while De Vos *et al.,* 2007 showed that only *ATG8e* was significantly induced in Arabidopsis upon 48 h and 72 h of aphid infestation [32]. In agreement with literature, here, we showed an upregulation of autophagy-related *ATG8-*family genes, *ATG8a*, *ATG8f*, and the cargo receptor *NBR1* upon 6 h of aphid infestation (Figure 1). Besides, a study reported that *ATG2-like, ATG6-like,* and *NBR1-like* genes were downregulated upon 6 h of *Rhophalosiphum padi* aphid infestation in a monocot plant, *Setaria viridis* [56]. The differences in the effect of aphid infestation on the *ATG* gene expression level might be related to the duration of infestation, and plant species. Another indication that the autophagy machinery is affected by aphid feeding is the increase in the number of autophagosomes. Many studies have shown that the number of autophagosomes in the cells is increased upon stresses such as fungus or virus pathogens [57,58]. A higher number of autophagic bodies was observed in the leaf tissues of GPA-infested Arabidopsis plants (Figure 2). Based on these results, we suggest that autophagy is induced in plants by phloem-feeding insects such as aphids.

### 3.2. Autophagy affects aphid performance and behavior

The *atg* mutants of Arabidopsis are generally described as being hypersensitive to abiotic stresses such as salt, osmotic stresses, and carbon starvation, as well as having leaf yellowing phenotypes and necrotic spots [40,59]. Studies have shown that the *atg* mutants are more susceptible to fungal necrotrophic pathogens [57]. Aphids that fed on *atg5.1* and *atg7.2* possessed lower body weight and poor fecundity relative to wildtype plants, indicating that *atg* mutants are more resistant to aphids (Figure 3). However, autophagy induction by pathogen attack has been shown to lead to different outcomes, either beneficial or detrimental for the host, depending on the pathogen’s lifestyle in plants. Studies have also shown that viruses could manipulate or hijack plant autophagy to modify nutrient availability to their benefit [21,60,61].

In addition, autophagy mutants were reported to have higher ROS levels, which might disturb GPA feeding [55,62,63]. Arabidopsis leaves produce ROS as a redox response to GPA infestation, and rapid ROS induction is often correlated with aphid resistance [64,65]. Thus, basal hydrogen peroxide levels in the mutants (*atg5.1* and *atg7.2*) were determined in the study, and a higher content was observed in *atg* mutants than in wildtype (Figure 7), which is consistent with previous studies that showed high accumulation of ROS in *atg* mutants [40,66,67]. Upon GPA feeding, hydrogen peroxide was induced in wildtype leaves, while levels in the mutants were higher than in the wildtype, and remained similar compared to untreated levels (Figure 7). We suggest that the high level of hydrogen peroxide might have caused a reduction in aphid feeding and reproduction in the mutants (Figure 3). It was previously reported that ROS are able to induce autophagy, while autophagy was also able to reduce ROS production [68]. Thus, the ROS induced by GPA feeding might trigger autophagy and the triggered autophagy might reduce ROS levels, which could be beneficial for GPA feeding [68–70]. We, therefore, suggest that autophagy-related mutations in Arabidopsis might cause either enhanced tolerance to insect attack or decreased attractiveness to insects in terms of phloem sap composition. In parallel, aphids might exploit the autophagy machinery to enhance their performance because the induced autophagy could reduce the plant’s defense mechanism against GPA stress *via* ROS production. In addition, the EPG analysis of aphids fed on *atg7.2* mutant plants showed poor feeding behavior, expressed in less feeding time in the phloem and more time in the epidermis and mesophyll tissues than wildtype, suggesting that GPAs were unable to acquire sufficient nutrients from the phloem sap. Notably, aphids fed on *atg5.1* plants showed a similar response as wildtype plants. Overall, the results suggested a difference in the composition of the mutant plants’ phloem sap. We, therefore, investigated the central metabolism of the *atg* mutants under GPA feeding.

### 3.3. Aphid feeding modified the phloem sap composition of *autophagy-deficient* mutants

Plants produce constitutive and inducible defensive compounds to protect themselves against insect attack while preserving their fitness [71]. The phloem sap of a host plant provides a carbon and nitrogen source for the invading insects [72]. It is known that the invading GPAs cause changes in the central metabolism of plants, such as carbohydrates and amino acids [73]. Carbohydrates are a major source of stored energy for host plants and insect herbivores, and amino acids are both growth-limiting for insect herbivores and serve as precursors for many defense-related plant metabolites [73]. In this study, GPA feeding affected the quantities of amino acids, particularly serine, threonine, and valine. In agreement with Avin-Wittenberg *et al.* (2015), which showed a significant reduction of amino acids in *atg* mutants under carbon starvation, here we showed the levels of serine, threonine, and valine in *atg7.2* mutants were not affected by GPA feeding [74]. By contrast, compared to wildtype, the *atg5.1* mutant exhibited high levels of these compounds in response to GPA feeding, suggesting that ATG5-and ATG7-dependent autophagy are differentially affected by aphids. These differences might be because both enzymes belong to two conjugation systems for autophagosome formation [75].

Furthermore, a unique set of central metabolites was presented that were altered in the *atg* mutants upon GPA feeding. Of these, amino acids and sugars were highly accumulated in wildtype and *atg5.1* mutants, while organic acids were increased in *atg7.2* mutants upon GPA feeding (Supplementary Figure S2). Wu *et al.* (2020) showed that restriction of dietary amino acids decreased the body weight of GPAs [76]. However, the effect of phloem sap composition on aphid performance or feeding behavior is more complex than a simple correlation with the nitrogen content of the diet. To conclude, the results of central metabolism in the phloem of the mutants might explain the poor feeding behavior and performance of GPAs.

### 3.4. Conclusions

In this study, we show that autophagy is induced by phloem sap-feeding aphids in plants, as illustrated in Figure 8. Although GPAs showed poor feeding behavior and performance on the *atg* mutants, the defense mechanism of plants against GPAs via *PAD4* in the mutants was not functioning as fully as in the wildtype plants. This might partially be explained by the different phloem sap composition in the mutants. However, the high hydrogen peroxide phenotypes of the *atg* mutants could explain this observation [40,55]. Moreover, a high level of sugars and a lower level of ROS in wildtype might explain the fact that aphids showed better performance and feeding behavior even though the defense mechanism via SA signaling was activated. In agreement with autophagy’s proposed dual role in plant-virus interactions [60], we could assume that GPAs might be exploiting the autophagy machinery for their benefit to obtain nutrients such as sugars or reduce the plant’s defense mechanism via ROS accumulation. Nevertheless, the role of autophagy in the plant’s defense against insects requires further investigation.

**Figure 8.**
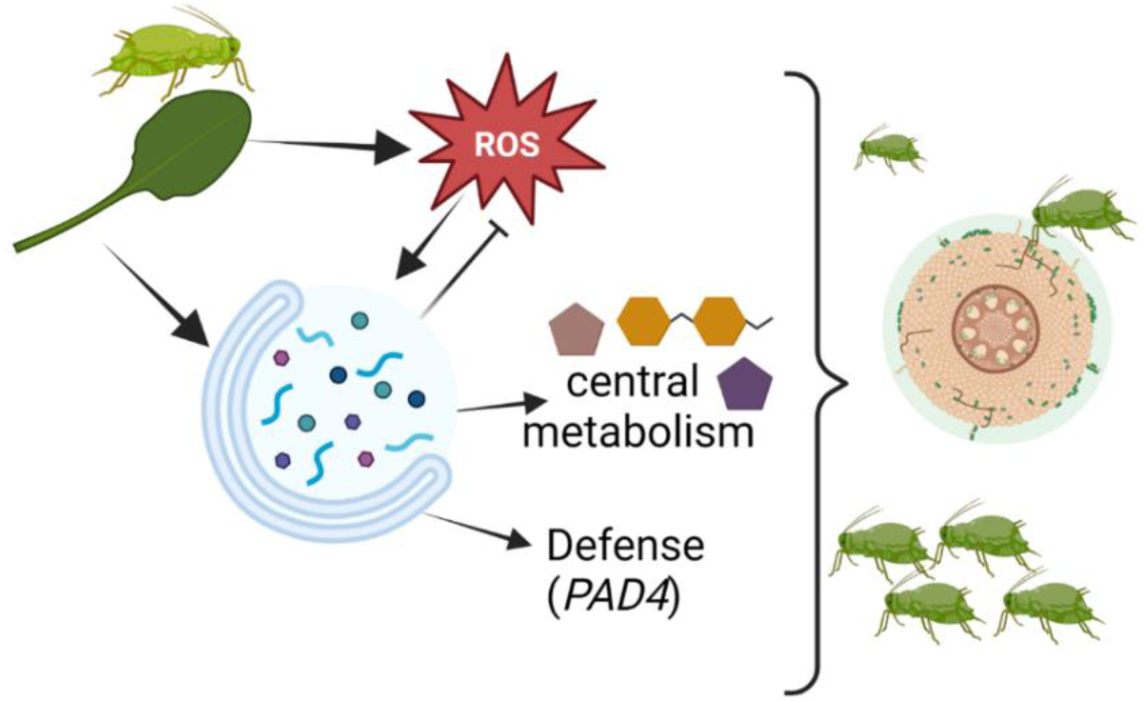
Proposed model of the autophagy mechanism under GPA infestation in Arabidopsis. Under GPA attack, autophagy-related genes or proteins are upregulated – such that autophagy is induced by aphid-induced stress in plants. The defense-related genes are also overexpressed – activating the plant’s defense against GPAs.

## 4. Materials and Methods

### 4.1. Plant material and growth conditions

*Arabidopsis thaliana* seeds were surface sterilized in 50% commercial bleach for 10 min to prevent the growth of microbial contaminants present on the seed surface and then rinsed three times with distilled water for 10 min [77]. The seeds were cold stratified at 4 °C in the dark for 4 d, then transplanted to 7 × 7 × 8 cm plastic pots filled with autoclaved Garden mix soil (70% peat, 30% perlite, fertilizer) and grown in a growth chamber with a photoperiod of 16 h light/ 8 h dark (120 μmol photons s^-1^ m^-2^) at 22 ± 3 °C. The *Arabidopsis thaliana* ecotype Columbia (Col-0) was used in this study. The Arabidopsis T-DNA insertion lines *atg5.1* (SAIL_129B079) and *atg7.2* (GK-655B06) and the transgenic line expressing GFP-ATG8f were previously described [62, 63, 64].

### 4.2. Aphid colony and bioassays

A green peach aphid (GPA; *Myzus persicae*) colony was provided by Prof. Shai Morin from Hebrew University of Jerusalem (HUJI), Israel, and reared on Arabidopsis Col-0 wildtype plants in a BugDorm (MegaView Science Co., Ltd., Taiwan) insect rearing tent (60 × 60 × 60 cm) with 96 × 26 μm mesh size. During the experiments, the GPAs were provided with the same environmental conditions as the plants (see above). For gene expression and metabolic profiling, 20 GPAs were confined to one rosette leaf of 4-week-old plants in a clip-cage (4.5 cm in diameter) for 6 h. For hydrogen peroxide detection, leaves were treated with GPAs for 7 d. As a control, the same setup was used, but aphids were not added into the clip-cages. Plant samples were then harvested, flash-frozen in liquid nitrogen and stored at -80 °C until further analysis. GPA body weight and fecundity measurements were conducted following Nalam *et al.* 2020 [78]. In brief, 20 adult GPAs were confined to a single leaf of each Arabidopsis genotype (wildtype, *atg5.1*, or *atg7.2*) within a clip-cage for 6 h. Subsequently, the GPAs were collected and weighed immediately using an analytical balance with a resolution of 0.01 mg (Satorius, Germany) to estimate body water content and body weight changes. Dry weights of the GPAs were obtained after drying the aphids at 55 °C for 8 h. Six biological replicates were used and independently repeated three times for each plant genotype in this experiment. For the fecundity experiments, aphids were synchronized by growing 50 adults on a Col-0 wildtype plant for 24 h. The new one-day nymphs (1^st^ instar) were allowed to reach adulthood (7 days). One of these adults was then confined to a single leaf of the different Arabidopsis lines, and the number of progeny was counted after seven days. Twelve biological replicates were used for each plant genotype in this experiment and independently repeated twice.

### 4.3. RNA extraction and qRT-PCR measurements

Total RNA was extracted using Sigma TRI-reagent (T9424) following the manufacturer’s protocol, then treated with DNase I to remove possible contamination of genomic DNA. The RNA concentration was quantified, and first-strand cDNA was synthesized with qScript™ cDNA synthesis kit (QuantaBio) from 1.5 µg of total RNA according to the manufacturer’s protocol. The integrity of newly synthesized cDNA was evaluated on a 2% agarose gel. The quantitative PCR reaction was performed using Power SYBR^®^ Green PCR Master Mix (Applied Biosystems, Foster City, CA, USA), according to the manufacturer’s protocol. Primers were designed using Primer-BLAST [79,80]. The accumulation of the target genes was normalized to the reference gene *Type 2A serine/threonine protein phosphatase* (*PP2A*) [81], for correction of technical variation in template amounts. Each sample was run in triplicates of the four biological replicates. The primers used for the qRT-PCR analysis are described in **Supplementary Table S1**.

### 4.4. Confocal imaging

A single leaf of the GFP-ATG8f transgenic plants was infested with 20 GPAs for 72 h. GPA-treated leaves of GFP-ATG8f plants were then incubated in 10 mM MES-NaOH (pH 5.5) buffer in the presence of 1 µM concanamycin A for 6-12 h in darkness at 23 °C. As controls, the same number of non-infested leaves were incubated in the incubation buffer with either dimethyl sulfoxide (DMSO) or with concanamycin A. An LSM 900 confocal laser scanning microscopy system (LSM 900, Zeiss, Germany) was used in this study. Generally, thin-section leaf samples were put between two microscope glass coverslips (No.1 thickness) in an aqueous environment. For image acquisition a Plan-Apochromat 40x/1.3 Oil DIC (UV) VIS-IR M27 objective was used on the Axio Imager.Z2 microscope GFP fluorescence images were taken using 488 nm laser excitation, and the emission was detected in the 490-550 nm range. The chlorophyll autofluorescence was imaged using the 638 nm laser and detected in the 645-700 nm range. Z-stack images composed of 20 to 50 images were taken using Z-stack, and snap images. The size of the recorded images was 159.73 × 159.73 µm (1744 × 1744 pixels). The pinhole diameter was 40 µm on all recordings. All acquired images were converted to CZI and TIFF formats using the Zen 3.1 (blue edition) image processing software. The experiment was conducted with six biological replicates, and the GPA-treated leaves of GFP-ATG8f plants were sectioned into four pieces as technical replicates.

### 4.5. Electrical Penetration Graph (EPG) analysis

GPA feeding behavior was monitored on wildtype and the two *atg* mutants, *atg5.1* and *atg7.2*, using the EPG on a GIGA 8 complete system (EPG Systems, Wageningen, the Netherlands) [82]. A dorsal surface of each adult GPA abdomen was attached with 18 μm diameter gold wire using silver glue [83]. One-month-old Arabidopsis plants were placed into a Faraday cage, electrodes were placed into the pots, then the aphids were allowed to contact the leaf surface, and their probing was adjusted. The GPAs were allowed to feed for 8 h, while the feeding behavior was recorded. For consistency with other experiments, only the first 6 h of the electrogram were analyzed. The waveforms were digitized at 100 Hz with an A/D converter, and patterns were recognized as described previously [82,84]. A computer was connected to the Giga direct current amplifier, and the waveforms were collected every 30 s with Stylet^+^d software (v01.30). The feeding behavior of GPAs on wildtype and *atg* mutants was compared by analyzing the time spent in each of the four main phases: pathway phase (PP), non-probing phase (NP), sieve element phase (SEP), and xylem phase (G). The subphases within SEP that indicate phloem salivation (E1) and phloem ingestion (E2) were also analyzed. Parameters such as the time to 1^st^ probe, the total number of probes, and the number of potential drops (PD) that indicate GPA health [85] were measured. The potential E2 index, number of E1 and E2 waveforms, total time spent in E1 and E2, and percent time spent in E2 greater than 10 min indicate phloem acceptability and plant defense response n [44]. EPG waveforms were analyzed using Stylet+a software and an Excel workbook for automatic parameter calculation as previously described [66, 95,104]. The experiment was repeated until 15 replicates were obtained for each treatment. However, a recording was not considered a replicate if GPAs spent more than 70% of the recording time in the non-probing, xylem, and derailed stylet phase. Thus, the final number of replicates for each treatment differed, i.e., wildtype = 13, *atg5.1* = 12, *atg7.2* = 14. The data were rank transformed, and differences between means were determined using ANOVA [87]. The proportions were compared using the Wilcoxon test with Steel’s method for nonparametric multiple comparisons with control.

### 4.6. Metabolite analysis using gas chromatography-mass spectrometry (GC-MS)

Approximately 100 mg of leaf homogenates were weighed in a 2 ml Eppendorf Safe-lock tube, and 1 ml of pre-cooled extraction mixture, methanol/methyl-tert-butyl-ether/water (1:3:1 v:v:v), was added to each tube and vortexed. Then, the samples were shaken on an orbital shaker at 1000 rpm at 4 °C for 10 minutes, followed by incubation in an ice-cooled ultrasonication bath for another 10 minutes. Next, the metabolites were phase-separated by adding 500 μl of UPLC-grade methanol/water (1:3 v:v). Samples were vigorously vortexed and centrifuged at 17,000xg at 4 °C for 7 min. The polar phase (200 μl) was transferred into a new tube, dried overnight in a SpeedVac (Thermo Scientific, USA) and stored at -80 °C [88]. Dried samples were derivatized before the GC-MS analysis. For derivatization, 40 μl of 20 mg methoxyamine hydrochloride (Sigma-Aldrich, UK) dissolved in 1 ml of pyridine was added to the dried sample and shaken on an orbital shaker at 1000 rpm at 37 °C for 2 h. Next, 70 μl of N-methyl-N-(trimethylsilyl) trifluoroacetamide (MSTFA) and 7 μl of alkane mix were added and shaken at 37 °C for 30 min. The derivatized sample (110 μl) was transferred to a vial and analyzed on a GC-MS machine. The mass spectrometry files were processed using the Agilent Mass Hunter software, and feature (mass peak) retention times and *m/z* were calculated. Annotation and quantification of detected metabolites were carried out with the Mass Hunter software, the NIST mass spectral library, and retention index (RI) libraries (gmd.mpimp-golm.mpg.de) [89]. Compounds were identified by comparing their retention index (RI) and mass spectrums, generated from authentic standards and libraries (Max-Planck Institute for Plant Physiology in Golm (http://gmd.mpimp-golm.mpg.de/) [88,90]. The metabolite response values were normalized to the internal standard, ribitol (Sigma-Aldrich, USA), and their respective tissue weights.

### 4.7. Detection of hydrogen peroxide

A 3,3′-diaminobenzidine (DAB) staining was used for *in situ* detection of hydrogen peroxide levels in wildtype and *atg* mutant plants [91]. GPA-treated or control leaves were gently vacuum-infiltrated with either DAB solution. As control, replicate leaves were infiltrated with buffer (10 mM Na_2_HPO_4)_. Samples were incubated in the DAB solution on a shaker for 4 h, then replaced with a bleaching solution (ethanol: acetic acid: glycerol (3:1:1)) to remove the chlorophyll and to visualize the precipitate formed by hydrogen peroxide (which renders precipitates in dark brown). Staining was done on three biological replicates for each treatment.

### 4.8. Statistical analysis

Student’s paired *t*-test and analysis of variance (ANOVA), were performed using Excel and JMP (SAS; www.jmp.com, USA) [92], respectively. Advanced Metaboanalyst 5.0 online software was used for metabolite analysis [93]. For Metaboanalyst analysis, the metabolite data were transformed into log_10_ values for normal distribution. For multiple testing analyses, *P*-values were adjusted according to Benjamini and Hochberg procedure (false discovery rate; FDR). Statistical significance was denoted when *P* values were less than 0.05, as indicated by an asterisk, respectively.

## Supplementary Materials

Figure S1: Validation of the expression level of *ATG* genes on the two *autophagy-defective* mutants used in the study; Table S1: Primers used for quantitative RT-PCR analysis; Table S2: Feeding behavior of GPAs on *atg* mutants; Table S3: Central metabolites detected in the phloem of Arabidopsis *atg* mutants and wildtype under GPA feeding; Table S4: Fold change values of significant metabolites affected either by one of the treatments or both.

## Author Contributions

Conceptualization, L.K.H., H.Z., and V.T.; methodology, L.K.H., R.S., A.D., S.M., W.J.P; validation, L.K.H., R.S., A.D., and S.M.; formal analysis, L.K.H., W.J.P., V.N., S.M.; investigation, L.K.H., S.M., and, W.J.P; resources, A.D., and Y.B.; data curation, L.K.H., writing—original draft preparation, L.K.H., H.Z., and V.T.; writing—review and editing, L.K.H., V.M., Y.B., S.M. H.Z. and V.T. All authors have read and agreed to the published version of the manuscript.

## Data Availability Statement

Data is contained within the article or Supplementary Material

## Acknowledgments

We are grateful to Noga Sikron Peres (BGU) for her assistance with the GC-MS and confocal microscopy, Valeria Mitsurova for the technical support and to Beery Yaakov for helping with primer design and RNA analysis. From the Hebrew University of Jerusalem, Israel, we thank Shai Morin for providing a *Myzus persicae* colony.

